# The Multivariate Normal Distribution Framework for Analyzing Association Studies

**DOI:** 10.1101/208199

**Authors:** Jose A. Lozano, Farhad Hormozdiari, Jong Wha (Joanne) Joo, Buhm Han, Eleazar Eskin

## Abstract

Genome-wide association studies (GWAS) have discovered thousands of variants involved in common human diseases. In these studies, frequencies of genetic variants are compared between a cohort of individuals with a disease (cases) and a cohort of healthy individuals (controls). Any variant that has a significantly different frequency between the two cohorts is considered an associated variant. A challenge in the analysis of GWAS studies is the fact that human population history causes nearby genetic variants in the genome to be correlated with each other. In this review, we demonstrate how to utilize the multivariate normal (MVN) distribution to explicitly take into account the correlation between genetic variants in a comprehensive framework for analysis of GWAS. We show how the MVN framework can be applied to perform association testing, correct for multiple hypothesis testing, estimate statistical power, and perform fine mapping and imputation.

## 1 Introduction

In the last decade, genome-wide association studies (GWAS) have discovered thousands of common variants implicated in genetic diseases [31]. Technological developments in microarray and sequencing technologies fueled these discoveries, which paved the way for cost-effective collection of genetic information in large amounts [32, 10]. Specifically, these technologies enabled the collection of genetic information at the scale of half a million variants spread throughout the genome. Further, the affordable cost of these technologies made feasible the study of thousands of individuals simultaneously. The initial GWAS studies [42] established essential groundwork for subsequent larger studies, which have identified the majority of today’s known common variants implicated in diseases [43].

While collecting genetic information on half a million variants is a technical marvel in itself, the actual amount of common variants present in the human genome is an order of magnitude larger. Fortunately, the correlation structure between genetic variants, referred to as “linkage-disequilibrium” (LD) in the genetics literature [36], makes the half million variants sufficient for GWAS [5]. The first large scale maps of human genetic variation [39, 13] partly aimed to identify this correlation structure. In fact, foundational literature enabling GWAS focused on identifying approaches to select the subset of variants that should be collected in a GWAS [5]. Even if a disease-causing variant is not collected in a GWAS, the locus (region of the genome) would be identified as associated given that a correlated variant is collected, which is referred to as a tag.

However, there are two sides of the coin with respect to LD. While the correlation structure of the human genome enabled important GWAS discoveries, the same correlation structure complicates GWAS analyses. Hypothesis tests of association at each variant are not independent due to this structure. Implications of LD include complicating multiple testing, complicating estimating statistical power, and introducing ambiguities to interpretation of association study results. This complication will only be exacerbated with the advance of next generation sequencing, which will enable future studies to collect virtually all of the genetic variants in the genome that are tightly correlated [38].

In this review, we describe a comprehensive approach to analyzing GWAS that uses the multivariate normal (MVN) distribution to model correlations in the genome. This approach offers an advantageous ability to model the effect of LD on all of the statistics simultaneously. The approach presented here provides a framework encompassing many different types of analyses related to GWAS, including multiple testing correction [3, 11, 19], estimation of statistical power [11], statistical fine mapping [16, 15, 24], and imputation [26, 34, 44] while taking into account the LD structure of the human genome.

## 2 GWAS at One SNP and Hypothesis Testing

### 2.1 Association Testing for Case/Control Studies

We first consider GWAS with case/control study design where information on genetic variants is collected from a dataset containing individuals with the disease (cases) and healthy individuals (controls). In this case, a hypothesis test is performed for each collected variant. This hypothesis test compares the frequency of a variant between the cases and controls in order to identify associated variants.

Here, we consider a GWAS study with a total of *n* individuals that are genotyped at *m* SNPs. In order to simplify notation, we assume a balanced case-control study where we have 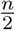 cases and 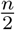 controls. Since each individual has two chromosomes, we have a total of *n* case chromosomes and *n* control chromosomes.

For each group and each SNP, we count the number of times that the minor allele appears and calculate the corresponding frequencies. Let 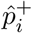 and 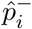 be the observed case and control frequencies, respectively, of SNP *i*. The true frequencies will be denoted as 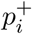 and 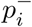. Assuming that *n* is large enough, the observed frequencies follow a Gaussian distribution 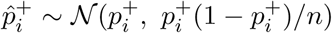 and 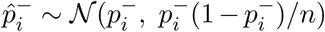 where 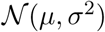 is a normal distribution with mean *µ* and variance *σ*^2^. We can then convert the observed frequencies into the following statistic

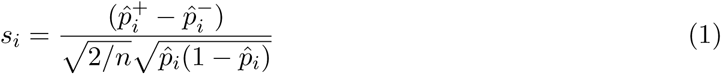

where 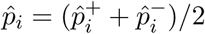. The statistic will follow the normal distribution

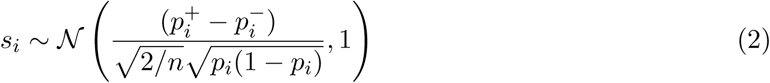

We denote the mean of this distribution as the non-centrality parameter 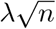 where

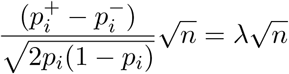

The probability density function of the normal distribution at point *x* for mean zero and variance *σ*^2^ is

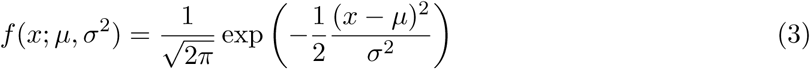

The statistic *s*_*i*_ takes into account the difference between the observed frequencies. When this difference is significantly high, we will assume an association between the SNP *i* and the disease. In GWAS, this is done under the framework of hypothesis testing. This framework allows us to control for Type I errors or quantify the error we can commit while implicating SNPs.

### 2.2 Association Testing for Continuous Phenotypes

The same framework can also be applied to continuous phenotypes such as cholesterol levels. We assume that our genetic study collects *n* individuals and the continuous phenotype of individual *j* is denoted as *y*_*j*_. We assume that the study collects *m* variants. We denote the frequency of variant *i* in the population as *p*_*i*_. We denote the genotype of the *i*th variant in the *j*th individual as *g*_*ij*_ *∈* {0, 1, 2}, which encodes the number of minor alleles for that variant present in the individual. In order to simplify the formulas and without loss of generality, we standardize the genotype values such that 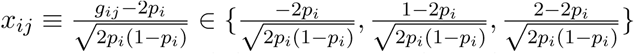 since the mean and variance of the column vector of genotype (*g*_*i*_) is 2*p*_*i*_ and 2*p*_*i*_(1 − *p*_*i*_), respectively. Due to the standardization, the sample mean and sample variance of the vector of genotypes at a specific variant *i* denoted as *X*_*i*_ are 0 and 1, respectively.

For the association at SNP *i*, the following model for the effect of SNP *i* on the phenotype is utilized

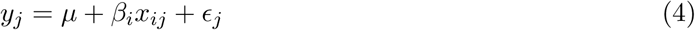

where *µ* is the population mean of the phenotype, *β*_*i*_ is the effect size of the SNP and 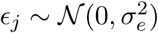 is the contribution of the environment to the phenotype for individual 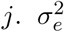 is referred to as the environmental variance. In vector notation, this model is

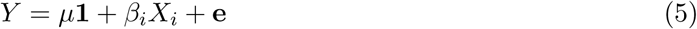

where *X*_*i*_ is a column vector of standardized genotypes for variant *i* and 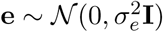, where **I** is the identity matrix of dimension *n*.

Using equation (5), we can obtain an estimate of *β*_*i*_ with the observed data. This reduces the equation to a simple regression problem where the resulting estimates are 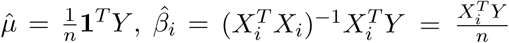 since *X*_*i*_ is standardized so 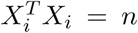. The estimated residuals 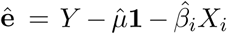 can be used to estimate the standard error 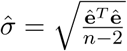. Since these studies are often quite large, the association statistic will approximately follow a normal distribution such that

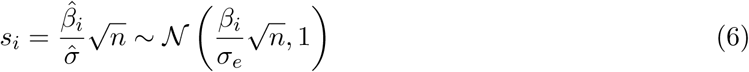

where the non-centrality parameter 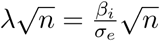.

### 2.3 Hypothesis Testing

If we assume that SNP *i* is not involved in the disease, referred to as the null hypothesis, then 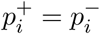, and therefore *s*_*i*_ follows a standard normal distribution 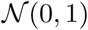. Typically, a false positive rate *α* (called type I error) is determined in advance (common values are 0.05 or 0.001 for a single hypothesis and 5 *×* 10^−8^ for a GWAS). From *α*, a threshold is calculated using the inverse of the standard normal cumulative distribution function Φ^−1^, i.e. *θ*_*α*_ = Φ^−1^(1 − *α*/2). A SNP *i* is declared as associated if |*s*_*i*_| > *θ*_*α*_. In this framework, *α* is the probability that *s*_*i*_ is in the tails of the standard Gaussian distribution under the assumption that the null hypothesis is true and the mean value of *s*_*i*_ is 0 (see Figure 1).

Often, a *p*-value is reported as the result of a statistical test. In this case, the *p*-value of a SNP *i* is the probability of observing *s*_*i*_ or a more extreme value assuming that SNP *i* is not associated. This value can be calculated using *p* = 2(1 − Φ^−1^(*s*_*i*_)) in case of *s*_*i*_ > 0 or *p* = 2Φ^−1^(*s*_*i*_) in case *s*_*i*_ < 0. Comparing *s*_*i*_ with the threshold is equivalent to comparing the *p*-value with *α*. If *p* < *α* then we declare the SNP as associated.

### 2.4 Statistical Power

When performing association testing, the null hypothesis assumes that the SNP is not associated with the disease. However, our objective is to discover the SNPs that are involved in the disease. In order to discover a SNP involved in the disease, when we perform the statistical test on collected data, we must reject the null hypothesis and declare the association as significant. We are interested in computing the probability of rejecting the null hypothesis for the SNPs which actually affect the disease. Intuitively, this is a measure of how likely an association study will succeed in finding the true disease causing variants. This quantity is referred to as the statistical power and can be calculated using the hypothesis testing framework. The power is a function of the effect size, which measures the effect of the SNP on the disease and the significance threshold. If SNP *i* is associated then 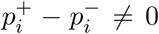 or *β*_*i*_ ≠ 0 and *s*_*i*_ is normally distributed with mean 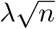 (the non-centrality
parameter) which is non-zero, and unit variance.

In order to declare a SNP *i* as associated, it has to happen that *|s_i_|* > *θ*_*α*_, so to know the probability of detecting it we have to calculate *Pr*(*|s_i_|* > *θ*_*α*_). However, in contrast with the case where SNP *i* was not associated and *s*_*i*_ followed a standard normal distribution, now *s*_*i*_ follows the Gaussian distribution 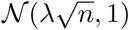. The power is visualized as the green area shown in the graphics presented in Figure 1. The power is a function of *α* and the non-centrality parameter 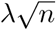 and can be computed using the following formula

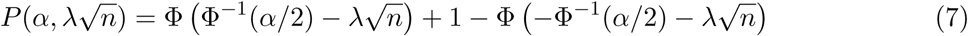

Implicitly, the power depends on factors such as the significance threshold, effect size, the minor allele frequency, and the number of individuals. Furthermore, a higher non-centrality parameter produces a higher power.

**Figure 1:**
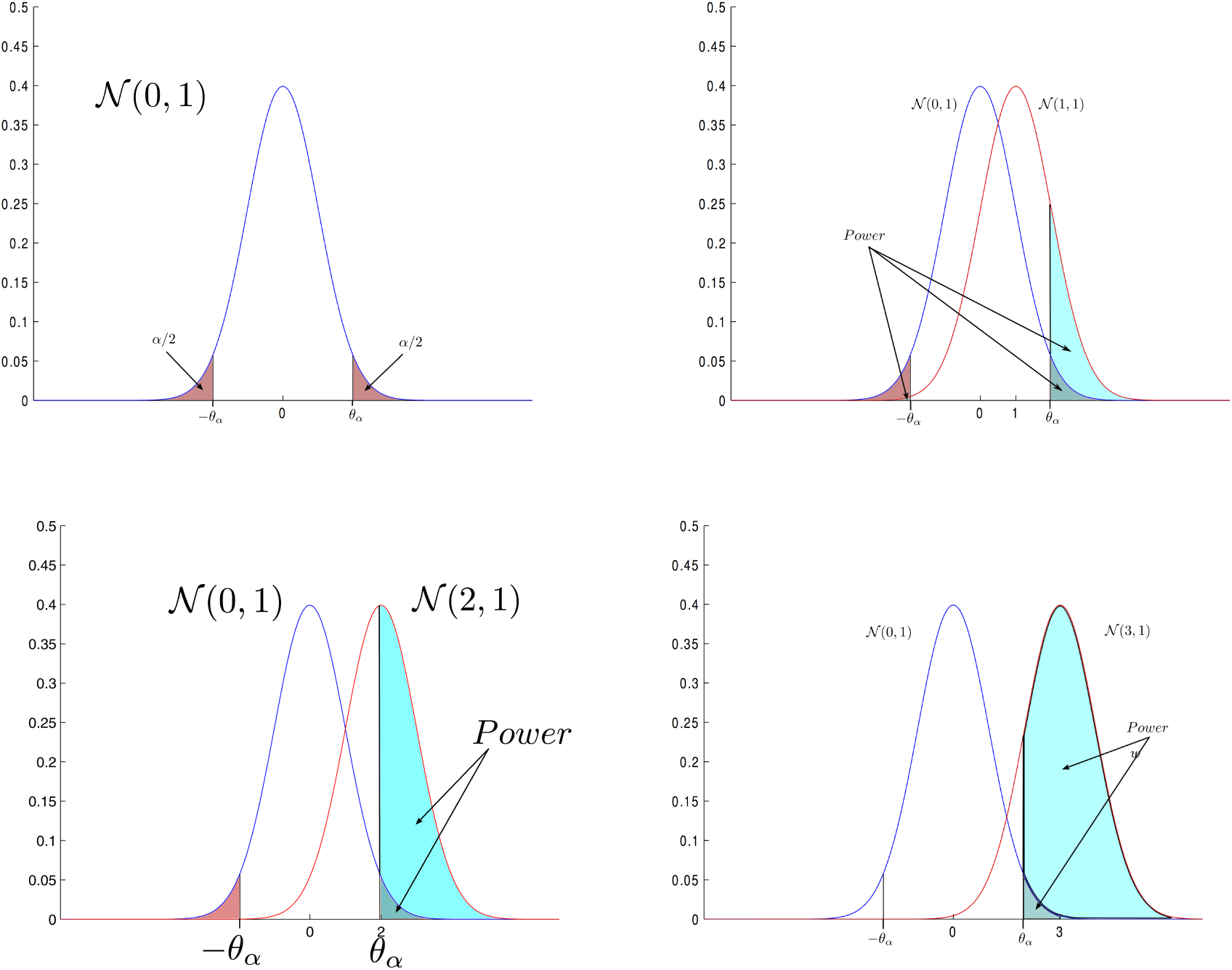
(a) A standard normal distribution with the reject region for an error *α*. (b) A 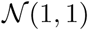 for an associated SNP with *λ* = 0.1 and *n* = 100. (c) A 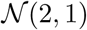 for an associated SNP where *λ* = 0.2 and *n* = 100. (d) A 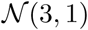 for an associated SNP where *λ* = 0.2 and *n* = 325

## 3 GWAS for Multiple SNPs

### 3.1 Correlated SNPs– Multivariate Normal Distribution Model

In a region of the genome that is involved in a disease, some of the variants will have a direct effect on the disease. We refer to these variants as causal variants. However, due to the correlation between the variants, many more of the variants will be associated and have non-zero non-centrality parameters. Association studies perform an association test at each SNP. Resulting statistics are dependent due to the underlying correlation structure, referred to as Linkage Disequilibrium (LD), of the SNPs themselves. A natural measure of the correlation between two SNPs (*i* and *j*) is simply their correlation coefficient, which can be calculated as follows

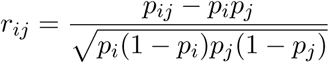

where *p*_*ij*_, *p*_*i*_, and *p*_*j*_ are the joint minor allele frequency of SNPs *i* and *j*, and the minor allele frequency of SNPs *i* and *j*, respectively.

Obviously, if two SNPs *i* and *j* are correlated then the probability distributions of the SNPs statistics, *s*_*i*_ and *s*_*j*_, should also be related. In fact, the correlation coefficient plays a central role in the joint distribution of the statistics of the *m* SNPs, *S*^*T*^ = (*s*_1_, *s*_2_, …, *s*_*m*_), which follows a multivariate normal distribution (MVN) [11, 25, 34, 15, 14, 17, 19, 44, 18]

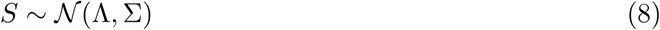

where 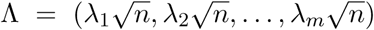 is the vector of non-centrality parameters and Σ is the variance-covariance matrix with 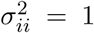 and *σ*_*ij*_ = *r*_*ij*_ for all *i* ≠ *j*. Σ is referred to as the LD matrix. The probability density function of the MVN at point *X* for mean vector *µ* and variance-covariance matrix Σ is

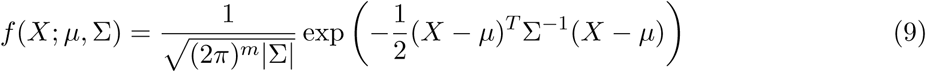

A derivation of the covariance is provided in Appendix A.

The non-centrality parameters for the SNPs depend on which SNPs are causal, the SNP effect sizes, and the correlation between the SNPs. If we consider two SNPs (*i* and *j*) where SNP *j* is a causal SNP with non-centrality parameter 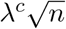, the non-centrality parameter of another SNP statistic 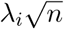 is

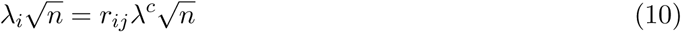

The previous equality has important implications for GWAS. Given that SNP *j* is a causal SNP, then 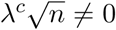 and, therefore, for each SNP *i* correlated with SNP *j* its non-centrality parameter 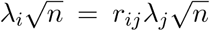 is also non-zero. This means that a GWAS study has a high probability of discovering both causal SNPs and highly correlated SNPs among the identified associated SNPs. Thus, not all SNPs must be collected in a GWAS for the purpose of identifying an associated region; only a subset of the SNPs (referred to as tag SNPs) need to be collected that are correlated with the remaining uncollected SNPs.

When we consider more than one SNP at a time in order to identify responsibility for the association, a distinction arises between the effect of the variants and the variants themselves. Observed effects of these variants can simply be due to the correlation between the effects and the causal variants. This distinction parallels the distinction between direct and indirect effects in the causal inference literature [35] and has been discussed at length in the genetics literature [36]. We use the notation 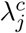 and the term effect size to denote the actual causal effect of SNP *j*. We note that the correlated SNP *i* has a non-centrality parameter 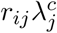 due to the correlation. We also note that if SNP *j* is causal, the actual non-centrality parameter at SNP *j* may differ from 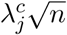 due to the effect of other variants in the region.

In general, where we can assume that the SNPs *j*_1_, *j*_2_, …, *j*_*k*_ are causal with individual effect size 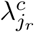 for *r* = 1, …, *k*, we can consider a vector *C*^*T*^ = (*c*_1_, *c*_2_, …, *c*_*m*_) such that

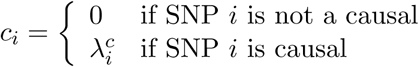

. Accounting for all this information, the multivariate normal can be written as

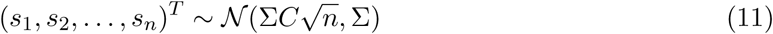

We note that 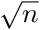 is scalar in the above equation that multiplies each entry in the mean vector, and we use the notation above for clarity. For example, suppose we have four SNPs where SNPs 2 and 4 are causal with effect sizes 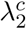 and 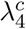 respectively. In this case vector *C* is as follows: 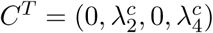 and the distribution of the statistics is

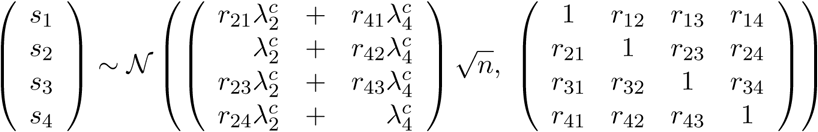

where we can see how the non-centrality parameter of each SNP is affected by the causal SNPs.

We note that many association studies today utilize linear mixed models for computing the assication statistics to take into account population structure or relatedness in the sample [21, 28, 29, 46, 45, 30]. When linear mixed models are utilized, the correlation between statistics is affected [20] and a derivation of the correlation when applying mixed models is shown in Appendix B.

### 3.2 Multiple Hypotheses Testing under the MVN Model

In a GWAS study, we carry out a hypothesis test for each SNP *i*. Although each of these hypotheses tests has associated a particular false positive rate *α*_*s*_, we would like to account for the overall false positive rate of the whole process of testing *m* non-associated SNPs. In the case of two SNPs *i* and *j*, the false positive rate can be viewed as the probability under the distribution

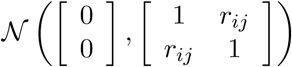

of the external part of a rectangle defined by the points 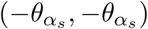 and 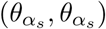 where 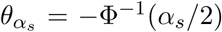 is the per-maker threshold associated to the SNPs (Figure 2). In the case of multiple SNPs, this region, which is the external part of a hypercube, is denoted as 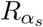. This region is the rejection region, where we reject at least one of the null hypotheses if our vector of statistics falls inside it. In the case of *m* SNPs, and a given per SNP threshold *α*_*s*_, the overall false positive rate of the study can be written as follows

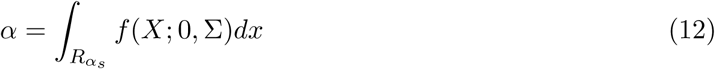

and the per SNP threshold *α*_*s*_ can be set so that the overall false positive rate of the study is at the desired level. From the equation it is clear that the false positive rate depends on the per SNP threshold, the number of SNPs and the variance-covariance matrix. When applied to a GWAS, equation (12) requires an integration in a space that may have over a million dimensions. An efficient method for computing this integration is described in Han et al., (2009)[11].

### 3.3 Power under the MVN Model

In order to calculate power under the MVN model, we assume that we know the vector of true effect sizes *C*. Therefore, the statistics (*s*_1_, *s*_2_, …, *s*_*m*_)^*T*^ follow a multivariate normal distribution 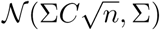 with *C* and Σ as in (11). Therefore, the power is the probability of the rejection region, 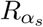, under the previously defined distribution. Specifically, the power then generalizes equation (7) to multiple SNPs

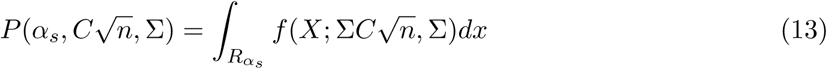

For example, in the case of two SNPs where the first SNP is the causal SNP with non-centrality parameter 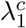, the correlation between the two SNPs is *r*_12_, and the per SNP significance threshold is *α*_*s*_, then the power of the association study testing both SNPs is

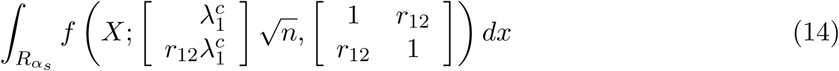

where 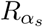 is the region defined outside of the square defined by the points 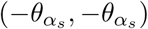 and 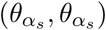. See Figure 2 for an example.

**Figure 2:**
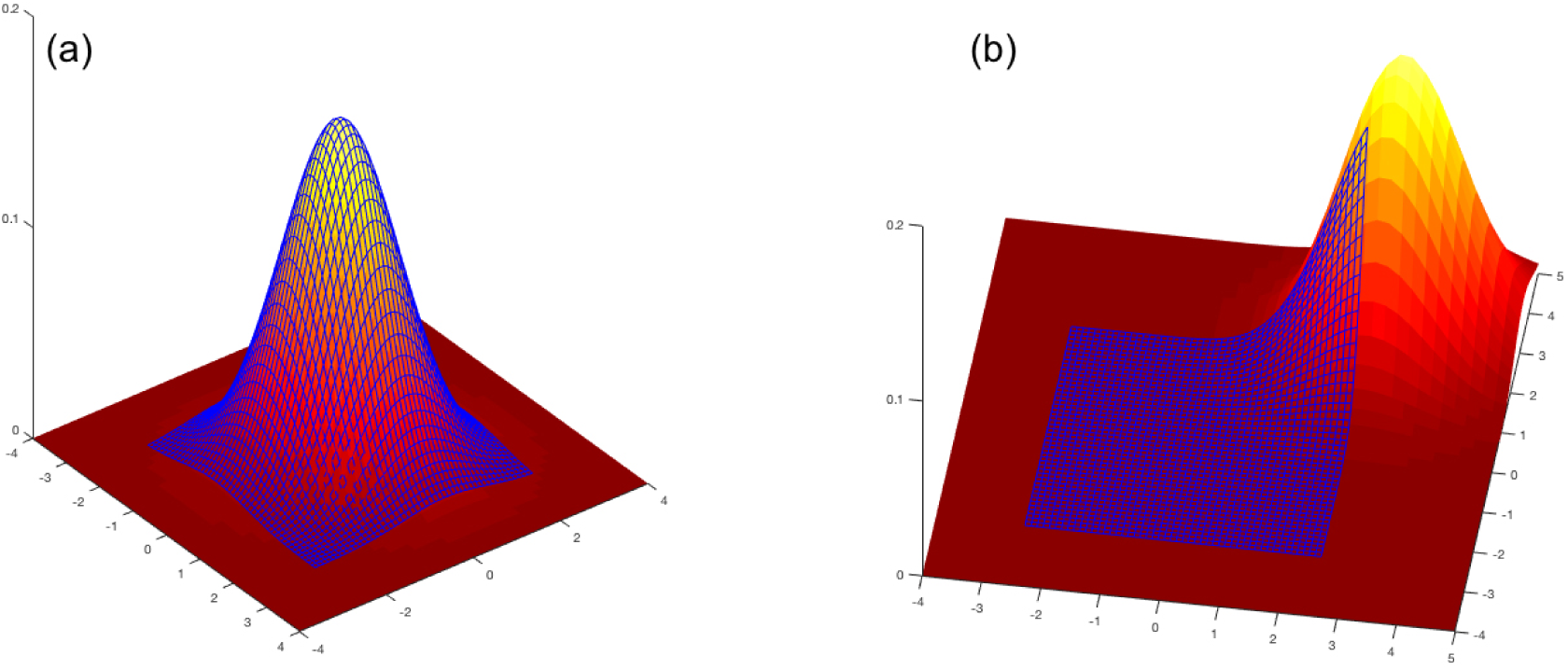
An illustration of the multivariate normal model (a) Type I Error (b) Power

The traditional notion of statistical power assumes a specific alternative hypothesis, which defines which variants are causal and makes an assumption on their effect sizes. However, in practice, we are interested in the concept of “average power” which is the average of the statistical power computed for each variant given a specific effect size.

In order to estimate the power of GWAS, we typically assume that each SNP has equal chance of being causal. We then compute the power for each SNP and report the average value over all the SNPs. This approach assumes a probability model over the causal vectors *C* where each SNP has equal chance of being the causal variant. A simple probability model is to assume that at most 1 SNP is causal and each SNP *i* has a probability of *c*_*i*_ (with 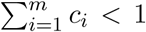) of being causal, all with the same effect size of *λ*^*c*^. In this scenario there are *m* + 1 possible models. We can define this set of possible models as 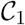. This set contains *m* + 1 vectors that can be denoted as *C*^(*i*)^ with *i* = 0, …, *m*, where *C*^(0)^ = (0, …, 0) represents the situation in which no SNP is causal (and has probability 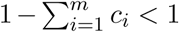) and *C*^(*i*)^, for *i* = 1, …, *m* is such that all the elements are 0 except the *i*th component that is *λ*^*c*^. The power of an association study in the case of using this probability model is

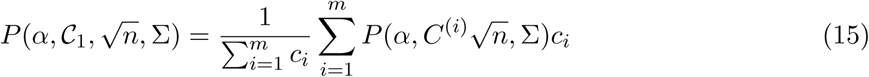

We note that each variant can be assigned a different prior probability based on additional data such as functional genomic data. We can then modify our association testing approach to maximize the statistical power in equation (15) as described in Eskin (2008) [7] and related publications [4, 6].

## 4 Statistical Fine Mapping

A central problem in GWAS is identifying the actual causal variants responsible for the association. This approach is referred to as statistical fine mapping. In this problem, we observe the results of an association study represented as a vector of observed statistics *S*, and we aim to obtain some information about the actual causal vector *C*. This requires assuming a distribution over possible values of *C*. We then define the prior over possible values of *C* as *P* (*C*). Given the value of *C*, we can then define the probably over the statistics as decribed above and we denote this as *P* (*S|C*).

The simplest formulation of fine mapping is to consider the set of probability models in 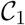. In this case, the posterior for each model *C*^(*i*)^, given the value of the statistics 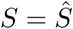, is

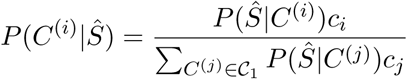

It can be noted that the ranking of 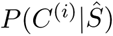 is the same as the one provided by the *p*-values of association statistics.

We can use these posteriors to define a “confidence set”. A confidence set is the set of possible causal variants that capture a sufficient fraction of the posterior distribution. Intuitively, this is the set of variants that has a high probability of containing the causal variant. We define the posterior for a set of *k* SNPs *j*_1_, *j*_2_, …, *j*_*k*_ as the sum of the posteriors 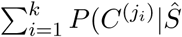. A typical “confidence set” is defined as a set that accounts for a posterior probability higher than .95. A confidence set computed for a GWAS locus intuitively is the set of SNPs that, with high probability, contains the causal variants responsible for the association at the locus.

Since the posterior computation requires assumptions, the interpretation of the “confidence sets” assumes that the priors over the causal vector *C* are consistent with what is actually occurring at the locus. In this sense, the model for statistical fine mapping 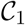 above is unrealistic for two reasons.

First, there are often multiple causal variants in the same locus, and our previous model only considers the scenario where one SNP affects the trait at a given locus. In general, this model is a good approximation if the causal variants are not in LD with each other. Secondly, it assumes that all the causal SNPs have the same effect size *λ*^*c*^.

Hormozdiari et al.,(2014)[16] introduced a hierarchical model based on the multivariate normal model which allows for multiple variants with effect sizes drawn from a normal distribution. Most recently developed fine mapping methods build upon this model[23, 2, 22, 1, 27]. In this case, we assume that each SNP has a probability *c*_*i*_ of being causal independently of the rest of SNPs. Here,
the set of models, denoted as 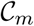 contains 2^*m*^ elements. If we define a binary variable *γ*_*i*_ that takes a value of 1 when SNP *i* is causal, and otherwise takes a value of 0, then the prior probability for a casual status is as follows

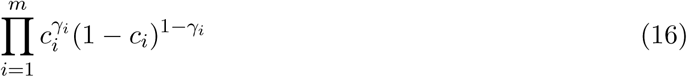

Once we know which SNPs are causal, we use a Gaussian model inspired by the classic Fisher’s polygenic model to get the effect sizes of the causal SNPs. The vector *C* is drawn from the distribution

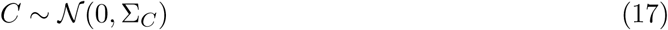

where Σ_*C*_ has elements

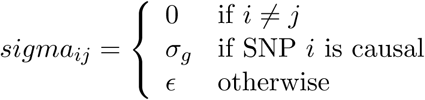

Under this model, the fine-mapping is carried out as follows. Given a set of SNPs 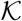, we denote the set of causal SNP configurations rendered by 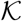 with 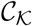, which accounts for all possible models with causal SNPs in 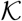. These are 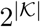 models, and the posterior probability of 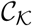 can be calculated as follows

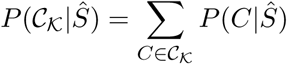

where to compute 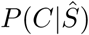, we utilize the Bayse’s rule and compute 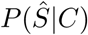 which is the likelihood of observed statistics given the vector of causal status. The details for this calculation is provided in Appendix C.

The fine-mapping consists in given a threshold *ρ* find the smallest subset of SNPs 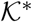 such that 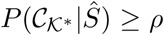. An algorithm to compute such a set is presented in Hormozdiari et al.,(2014) [16]. We can also incorporate functional genomics data to set the prior probabilities (*c*_*i*_) in the fine mapping model [24].

## 5 Inference about Uncollected SNPs

In the context of GWAS, one of the advantages of this multivariate framework is that it allows us to carry out inference about uncollected SNPs (i.e., given a non-collected SNP *u* we can use the statistics from correlated SNPs to obtain some information about the statistic of SNP *u*).

Given an non-collected or unobserved SNP *u*, we consider its *O* most strongly correlated collected SNPs (this information can be obtained from a database of SNPs such as the HapMap). Let *R*_*uO*_ denote the *O ×* 1 vector of the correlation coefficients between *u* and the *O* tag SNPs. Similarly, let *S*_*O*_ and *C*, respectively, be the *O ×* 1 vectors of the association statistics; let effect sizes of the tag SNPs and Σ_*O*_ be the *O × O* matrix of their pairwise correlation coefficients. The joint distribution of the association statistics of the unobserved SNP *u* and the *O* tag SNPs follows a multivariate normal distribution, which can be obtained by developing equation 11 and expressed as follows

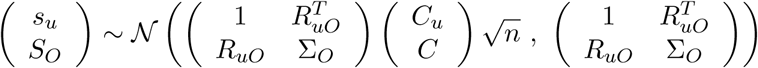

The previous joint distribution allows us to make inference about the statistic of the unobserved SNP *u*. Departing from that distribution, we can calculate the distribution of the statistic of the uncollected SNP *u* given the statistic of the collected SNPs 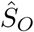

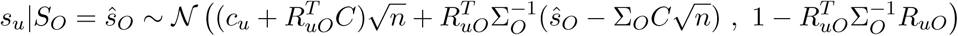

This distribution can be used in different ways. This approach can be thought of as a method for imputation [26, 34]. For example, we could fill the value of *s*_*u*_ with the mean value of the previous distribution. A second application is to calculate the probability of that SNP being causal given the value of the collected SNPs [25, 44].

## 6 Discussion

We have presented how the multivariate normal distribution can be utilized to explicitly model the correlation between variants in the analysis of GWAS studies. We have demonstrated how the MVN framework can be applied to correct for multiple testing, estimate the statistical power of an association study, and perform fine mapping and imputation.

## A Derivation of covariance of association statistics

In this section, we show the derivation of the covariance between association statistics. Let *m* be the number of SNPs, *s*_*i*_ be a statistic for the *i*th SNP, and Σ = {`Cov`(*s*_*i*_, *s*_*j*_)} be the *m × m* covariance matrix between the statistics.

For the association at SNP *i*, the following model for the effect of SNP *i* on the *k*th individual is utilized

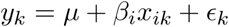

and in vector notation

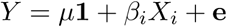

Here, *X*_*i*_ is a column vector of normalized genotypes for variant *i* and 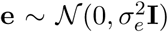, where **I** is the identity matrix of dimension *n* and **1** is a column vector of 1’s. Then, the phenotype follows a MVN with a mean and variance as follows:

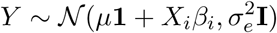

The ordinary least-squares solutions of *β* for SNP *i* and SNP *j* are as follows:

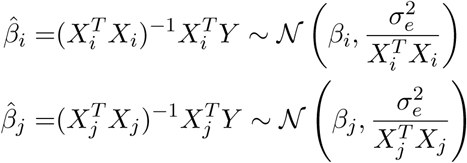

The association statistics of the two SNPs are computed as follows:

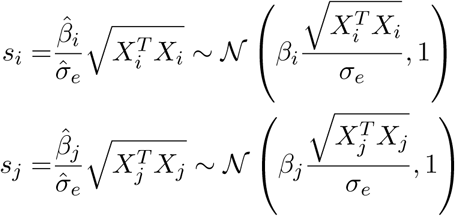

Here, the estimated values for *µ*, **e**, and *σ* for the SNP *i* are as follows: 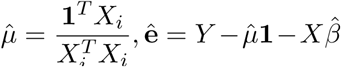 and 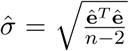. Then, we can prove that the covariance of the two statistics, `Cov`(*s*_*i*_, *s*_*j*_), is equal to the correlation between the genotypes, *r*_*ij*_, as follows:

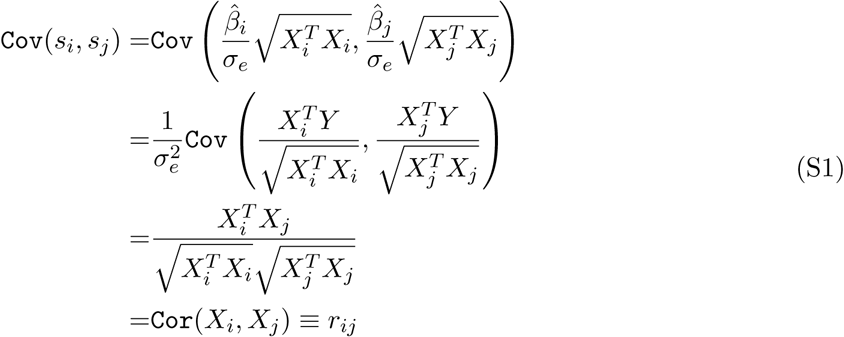

This relationship between genotype correlation and MVN covariance holds for case/control studies as well [37, 11].

## B Covariance of association statistics taking into account for population structure

Because of each population’s own genetic and social history, allele frequencies are known to vary widely from population to population. This creates genetic similarity between individuals in the study population, referred to as “population structure”. Individuals within a population have more similar phenotype values than individuals in distant populations. Population structure, along with this correlation of a phenotype with its populations, may cause spurious correlations between genotypes and a phenotype and induce an inflation of the values of association statistics leading to false positives [33, 9, 40, 12, 41, 8]. Linear mixed model (LMM) has emerged as a general approach to address this problem by explicitly modeling population structure in its association statistic [21, 28, 29, 46, 45, 30, 20].

For LMM, equation (S1) is no longer valid. That is, we cannot use the genotype correlation matrix as the covariance matrix of association statistics for mixed model. To derive the covariance matrix of association statistics under structured population, we assume a mixed model instead of the linear model equation shown in the previous section. For the association at SNP *i*, the following LMM for the effect of SNP *i* on the *k*th individual is utilized

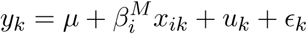

and in vector notation

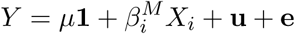

Here, *X*_*i*_ is a column vector of normalized genotypes for variant 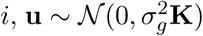 is a column vector modeling population structure effects, where **K** is the genetic relative matrix, referred to as “kinship matrix”, that explains the correlation between the individuals induced by population structure. Since the genotypes are normalized, the kinship matrix can be expressed as **K** = *XX*^*T*^ /*m*, where *m* is the number of genotypes and *X* is the *n×m* matrix of the normalized genotypes. 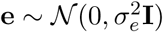 is a column vector modeling residual errors, where **I** is the identity matrix of dimension *n*.

Under this model, the phenotype follows a MVN with a mean and variance as follows:

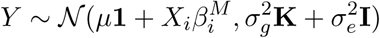

Given the observed data, it is straightforward to fit a LMM and estimate the parameters 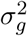 and 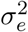 using standard strategies, which define the covariance matrix of phenotypes, 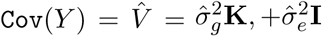. Now we utilize the fact that after obtaining 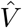, the remaining regression procedure is equivalent to performing ordinary least-squares in the transformed space,

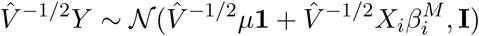

where both genotypes and phenotypes are transformed by a factor 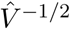. Assuming that 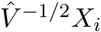 and 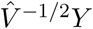 are normalized as mean 0 and variance 1 (without loss of generality), the ordinary least-squares solution of 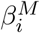 for *i*th SNP and *j*th SNP are as follows:

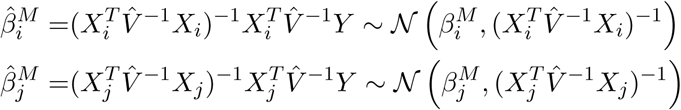

The statistics are computed as follows:

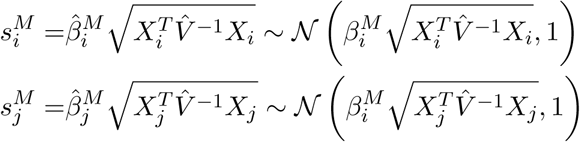

Accordingly, the correlation between the statistics changes from Equation (S1) to the following and the correlation between the statistics are equal to the correlation between the SNP transformed by the inverse square root of 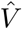,

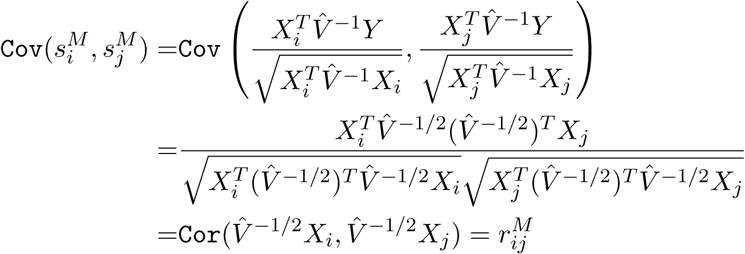

Thus, we can account for population structure in analyses related to GWAS, including multiple testing correction [3, 11, 19], estimation of statistical power [11], statistical fine mapping [16, 15, 24], and imputation [26, 34].

## C Efficient likelihood computation

We show in Equation (11) that the joint distribution of marginal statistics given the causal status is as follows:

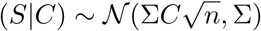

In addition, we use Equation (17) that is inspired by the classic Fisher’s polygenic model to get the effect sizes of the causal SNPs. This distribution is as follows:

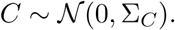

Utilizing the MVN conjugate prior and applying it to Equations (11) and (17), we can obtain the joint distribution of marginal statistics as follows:

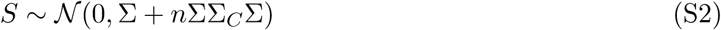

To compute the likelihood of the casual status, we utilize the probability density function of the MVN as shown in Equation (9). Unfortunately, a naive method to compute the likelihood is computationally intensive. In the naive method, we need to compute *S*^*T*^ (Σ + *n*ΣΣ_*C*_Σ)^−1^*S* and |Σ + *n*ΣΣ_*C*_Σ| that both require *O*(*m*^3^) operations. We use Woodbury matrix identity formula to speedup the computation of *S*^*T*^ (Σ + *n*ΣΣ_*C*_Σ)^−1^*S* and use Sylvester’s determinant identity to speedup the computation of |Σ + *n*ΣΣ_*C*_Σ|.

We reduce the time complexity by only computing the values that change with matrix *C*. We can factor out the Σ matrix as follows:

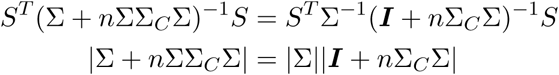

where |Σ| and *S*^*T*^Σ^−1^ can be computed once and can be used many times. Thus, we need to compute (***I*** + *n*Σ_*C*_Σ)^−1^ and |***I*** + *n*Σ_*C*_Σ| for every causal status. It is worth mentioning that we require Σ to be full rank. Unfortunately, in some loci, Σ, can be low rank. In this section, we assume that the LD matrix is full rank and in Appendix D we deal with low rank LD matrices. To ease the notation, we introduce two matrices *U* and *V* where *U* has (*m × k*) elements and *V* has (*k × m*) elements. We set elements of *U* and *V* such that *n*Σ_*C*_Σ = *UV*. Let *α*_*i*_ indicate the index of *i*th causal variant. We set elements of *V* as follows: 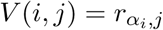. We set *U* (*α_i_, i*) to *nσ* while the rest of elements in *U* are set to zero.

We use the Woodbury matrix identity formula to compute (***I*** + *n*Σ_*C*_Σ)^−1^. The Woodbury matrix identity formula is as follows:

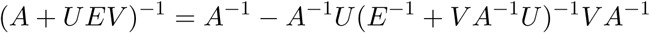

where we set *A* to ***I***_*m×m*_ and *E* to ***I***_*k×k*_. As a result, we have:

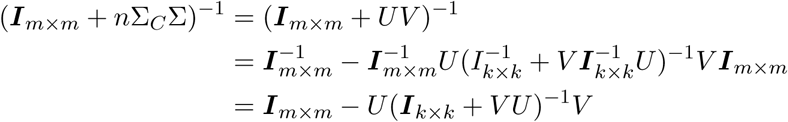

Interestingly, to compute (***I***_*k×k*_ + *V U*)^−1^ we need to inverse a *k × k* matrix that is much smaller than inverting a *m × m* matrix. Thus, we reduce the computation of *S*^*T*^ (Σ + *n*ΣΣ_*C*_Σ)^−1^*S* from *O*(*m*^3^) to *O*(*m*^2^*k*) where *k* is the number of causal variants for a given causal status (*k << m*).

We use the Sylvester’s determinant identity to speedup *|**I*** +*n*Σ_*C*_Σ| computation. The Sylvester’s determinant identity formula is as follows:

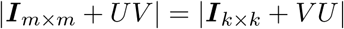

Thus, instead of computing the determinate of a *n × n* matrix, we can compute the determinate of a *k × k* matrix. We set matrices *U* and *V* such that *UV* = *n*Σ_*C*_Σ. Thus, we can compute *|**I*** + *n*Σ_*C*_Σ| in *O*(*k*^3^) operations [18].

## D Handling Low Rank LD Matrices

As mentioned in the above section, we assume that the LD matrix, Σ, is full rank. However, in some loci the LD matrix can be low rank due to linear dependency between different variants (e.g., two variants that are in perfect LD). We recall that we use Equation (S2) to compute the likelihood of each causal status. The LD matrix is computed from genotype data (Σ = *X^T^ X*), thus the LD matrix is semi-positive definite. Using the fact that the LD matrix is semi-positive definite, we can use the eigenvalue decomposition of the LD matrix which is as follows:

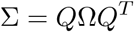

where *Q* is the matrix of eigenvectors and the *i*th column of *Q* is the *i*-th eigenvector of matrix Σ. Matrix *Q* is an orthogonal matrix (*Q^T^ Q* = *QQ*^*T*^ = ***I***). Let Ω be a diagonal matrix that consists of eigenvalues of Σ where the *i*th diagonal element of Ω is the *i*th eigenvalue of matrix Σ. We introduce a new set of marginal statistics *S′* = Ω^−1/2^*Q^T^ S* such that the joint distribution is computed as follows:

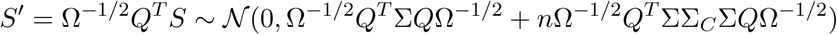

where we can simplify Ω^−1/2^*Q*^*T*^ Σ*Q*Ω^−1/2^ to ***I*** and *n*Ω^−1/2^*Q*^*T*^ ΣΣ_*C*_Σ*Q*Ω^−1/2^ to *n*Ω^1/2^*Q*^*T*^ Σ_*C*_*Q*Ω^1/2^ which can be shown as follows:

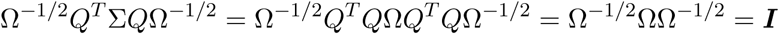

Similarly, we have:

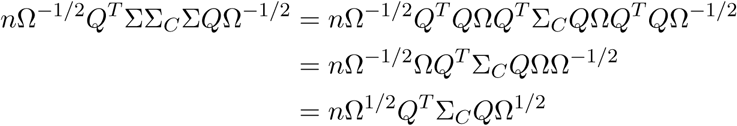

Thus, the joint distribution of *S′* is as follows:

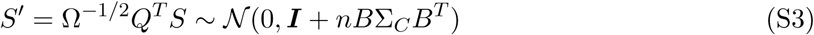

where *B* = Ω^1/2^*Q*^*T*^. It is worth mentioning that ***I*** + *nB*Σ_*C*_B^*T*^ is full rank. Thus, we can compute the likelihood of causal status for a locus where the LD matrix is not full rank.

